# Bacterial Growth Induced Biocementation Technology, ‘Space-Brick’ - A Proposal for Experiment at Microgravity and Planetary Environments

**DOI:** 10.1101/2020.01.22.914853

**Authors:** Aloke Kumar, Rashmi Dikshit, Nitin Gupta, Animesh Jain, Arjun Dey, Anuj Nandi, I. Venugopal, Koushik Viswanathan, N. Sridhara, A. Rajendra

**Affiliations:** Dept. of Mechanical Engineering, Indian Institute of Science, Bengaluru-560012, India.; U. R. Rao Satellite Centre, Indian Space Research Organisation, Old Airport Road, Bengaluru-560017

**Keywords:** Microbial induced calcite precipitation, Lunar simulant soil, space brick, bioconsolidation, space bricks

## Abstract

We present results of our investigation of microbial induced calcite precipitation for manufacturing ‘space bricks’ and a proposal for study of this activity in low-earth orbit (LEO). *Sporosarcina pasteurii*, a urease producing bacterial strain was used to consolidate lunar simulant soil (LSS) in the form of a ‘brick’ with non-trivial strength properties. Potential of a naturally occurring polymer namely, guar gum, as an additive was investigated for enhancement in compressive strength of bio-consolidated samples. Experimental results of bio-brick exhibited an approximate 10-fold increase in compressive strength with guar gum supplementation in soil. We present results of microstructural analysis of the ‘space bricks’ and also propose a payload design for related experiments in LEO.

## I Introduction

Around the world, space organizations are exploring technologies to develop extraterrestrial settlements [1]. As transportation of cement and other construction material from earth to the moon is economically daunting, there is a need to find an in-situ construction methodology using locally available material [2]. Ease of collection, potential radiation shielding properties, ability to withstand extreme lunar cycles and abundance make lunar regolith a promising material for the construction of infrastructure on the moon surface [3]. Moreover, lunar regolith has silty sand like characteristics with 46-110 μm grain sizes making it an ideal in situ resource [4].

Researchers have explored ways to establish in situ construction material manufacturing methodologies such as sintering using laser beams or microwave irradiation, cast basalt production, lunar glass preparation, thermite reaction, sulfur-based concrete preparation, dry-mix/steam injection method, 3D printing with different materials using lunar regolith [5]. Each method has its own sets of limitations. For instance, sintering has drawbacks such as high porosity, thermal cracking, difficulty in casting long blocks [6–8]. Nature has perfected the art of building strong and sustainable habitats such as corals, spider webs, termite mounds, and anthills with optimized resource consumption [8, 9]. Biomimicry might be a possible solution to make sustainable settlements on extra-terrestrial surfaces [10].

Microbial Induced Calcite Precipitation (MICP), a biomineralization process that uses the metabolic activities of calcifying bacteria has been explored as a method for bioconsolidation and self-healing structures [11, 12]. The urease pathway of MICP wherein ureolytic bacteria hydrolyse urea to make conditions favourable for mineral precipitation in short time has been widely explored [13–15]. Interestingly, Lambert and Randall [16] have shown that the green brick can be manufactured with the use of human urine as a urea source [17]. Researchers have also explored possible admixtures to strengthen the consolidated samples [18, 19]. Bio-based admixtures using additives such as xanthan gum, gellan gum, welan gum lignosulfonates are being explored by researchers for improvement in durability [20]. With a strong contemporary emphasis on bio-degradable, renewable and non-toxic material synthesis, there is an interest in the use of natural polysaccharides. Along with their biodegradable nature, naturally occurring polysaccharides present an economically viable material for manufacturing.

The aim of present work is to explore the utility of specific bacterial strains for manufacturing bricks using MICP and to identify potential routes for increasing the compressive strength of the resulting bio-consolidated bricks. We also briefly discuss future possibilities of conducting in situ consolidation experiments using miniaturized payloads under microgravity conditions. It is believed that this will help evaluate the practical utility of MICP as a long-term solution for building extra-terrestrial settlements.

## II Material and Methods

### A. Microorganisms and culture conditions

Bacterial-induced bio-consolidation was explored using ureolytic bacterial strains; namely, Sporosarcina pasteurii (Miquel) Yoon et al. ATCC®11859™, procured from American Type Culture Collection (ATCC).

### B. Lunar soil simulant

A new lunar soil simulant – LSS, developed and characterized by the Indian Space Research Organisation (ISRO) was used for the study [21]. Particle size characteristics of LSS are d10 (10% of sand particles have this diameter or lower) = 10 μm; d50 = 60 μm; d90 = 130 μm. Figure 1 represents the scanning electron microscopy (SEM) image of segregated mass of LSS and a & b represents elemental distribution calcium in LSS. Since the present work highlights the CaCO3 mediated bio-consolidation via bacterial activity, the presence and amount of calcium is important. LSS contains approximately 17.24 percent calcium in the form of CaO. Figure 2 c represents the spectrum of elements present in the LSS. Presence of Au was also identified by the spectrum, due to the gold coating performed to avoid charging of particles before SEM analysis.

**Figure 1.**
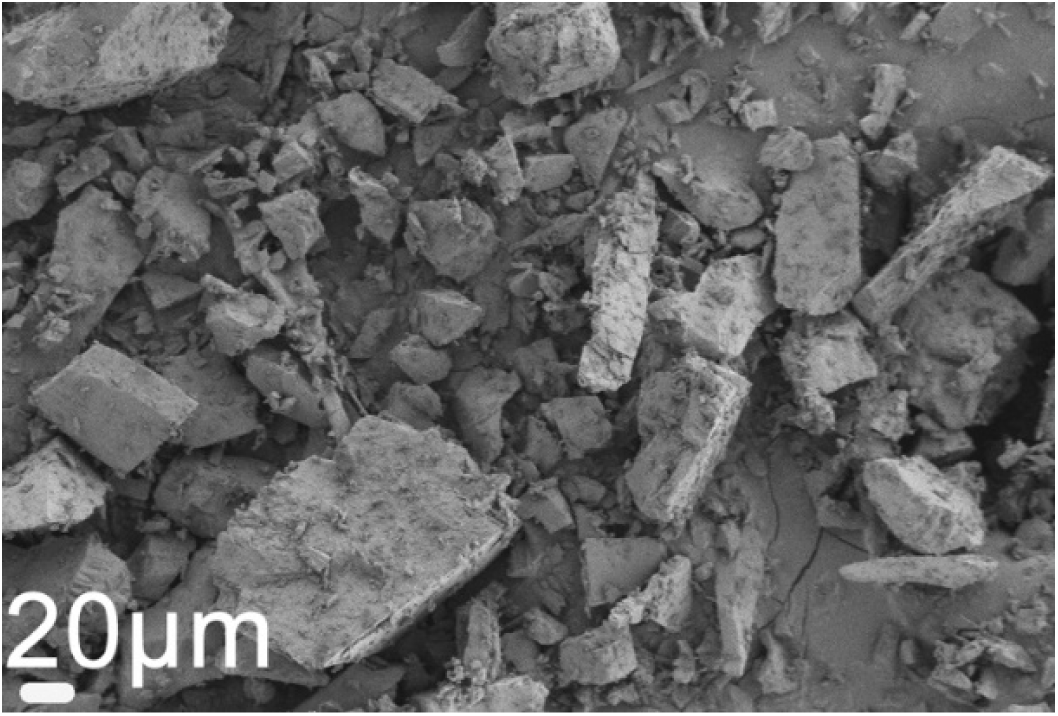
SEM image of loose mass of LSS

**Figure 2.**
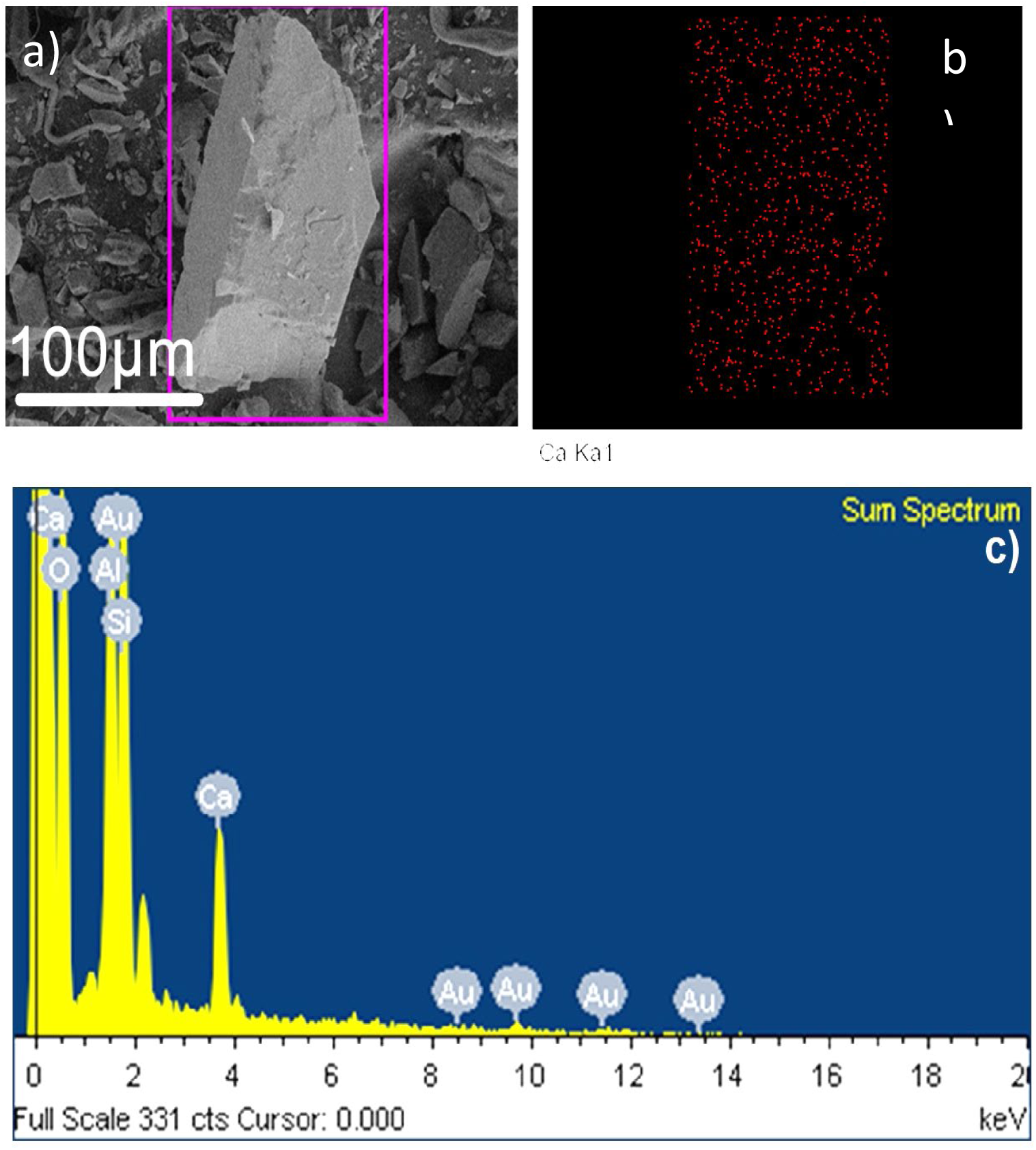
a) SEM image of LSS mapped location b) calcium distribution in LSS loose mass c) spectrum of elements present in LSS

### C. Preparation of sample for bio-consolidation and design of the bioreactor

A syringe column bioreactor was designed using 5 ml disposable polypropylene syringes (DispoVanTM) with an internal diameter of 10 mm. The outlet of the bioreactor was sealed with parafilm, and inner surface was layered with OHP transparency film to ensure easy removal of samples. The column was fitted with scouring pad at the bottom and packed with soil. The top portion of the bioreactor was kept empty for aeration and was covered with a cotton plug to avoid contamination. To avoid external contamination, LSS was autoclaved at 121°C and 15 psi for 30 minutes. Three sets of experiments were performed (without bacterium) with different concentrations of guar gum (GG) (0%, 0.5% and 1% w/w) namely, GG00LSSc, GG05LSSc, and GG01LSSc and were used as control. Since the ureolytic pathway of bacterial mediated consolidation was used in the present study, one set of experiment, namely, URGG00LSS, was also performed to check the role synthetic urease enzyme on the consolidation of LSS.

The media used (Synthetic Media-urea; SMU, hereafter) was prepared by adding 0.1 g peptone, 0.1 g glucose, 0.2 g mono-potassium phosphate, 0.5 g NaCl and 2 g urea in 100 grams of LSS. The nomenclature adopted along with the composition for various combinations is provided in Table 1.

**Table 1.** Different combination of treatments used in experiment for LSS consolidation

| Name | Bacteria | Media | Calcium lactate | Chemical urease | Guar Gum percentage |
| --- | --- | --- | --- | --- | --- |
| GG00LSSc | NA | SMU | Yes | No | 0% |
| GG05LSSc | NA | SMU | Yes | No | 0.5% |
| GG01LSSc | NA | SMU | Yes | No | 1% |
| SPSMULSS | SP | SMU | Yes | No | 0% |
| URGG00LSS | NA | SMU | Yes | Yes | 0% |
| SPGG00LSS | SP | SMU | Yes | No | 0% |
| SPGG05LSS | SP | SMU | Yes | No | 0.5% |
| SPGG01LSS | SP | SMU | Yes | No | 1% |

Urease enzyme (1 mg/g of LSS) extracted from *Canavalia ensiformis* (jack bean) was used in the present study and was procured from Sigma Aldrich, India. An experiment namely SPSMULSS was performed to specifically check the ability of bacterial strain or their secreted bio-products to bind the loose mass without the need of an external calcium source. Each experiment was performed in five sets. All chemicals were procured from Hi-media, India.

The bioreactor was filled with 7 g of LSS and the combination of treatments as provided in Table 2. 1.0 ml bacterial culture with optical density (OD600) of 1.0 was poured through the inlet of bioreactor. After 14 days of incubation in the bioreactor, fluid was drained out from outlet; samples were removed and kept for drying in a hot air oven (BioBee, India) at 45°C for a period of 12hr. Each experiment was performed in five sets. All chemicals were procured from Hi-media, India.

### D. Compressive strength measurement

Samples were prepared for compressive strength measurements with height to diameter ratio (H:D) between 1.8 and 2.2 as per ASTM C 39-04a. The surface of the consolidated samples was made flat by polishing with sandpaper of various grades (grit 2000). Compressive strength of consolidated samples was measured using a micro universal testing machine (UTM, Mecmesin, UK) under quasi-static conditions and a loading rate of 5 mm/min. A load cell of 500N capacity was used for force measurements. Displacement data was obtained using long travel extensometer (LTE) 700 designed by Mecmesin test system. For each case, five sets of samples were prepared, and the mean value was taken to be the representative compressive strength.

### E. Microstructural characterization

The microstructures of all consolidated bricks were examined under scanning electron microscope (SEM: Carl Zeiss AG – ULTRA 55). The relevant samples were collected and then air dried and ground to powder. The powder was then investigated by X-ray diffraction (XRD: powder XRD, Rigaku SmartLab, Rigaku Corporation) technique. Further thorough indexing of the XRD pattern was carried out by ICSD data based PANalytical X’Pert HighScore Plus software.

## III Result and Discussion

### A. Compressive strength of consolidated bricks

Potential of Sporosarcina pasteurii to consolidate LSS via MICP was investigated and it was found that the experimented bacterial strain was efficient enough to consolidate LSS. Figure 3 shows consolidated LSS sample after 14 days of incubation using the described bioreactor and inoculation with *S. pasteurii*.

**Figure 3.**
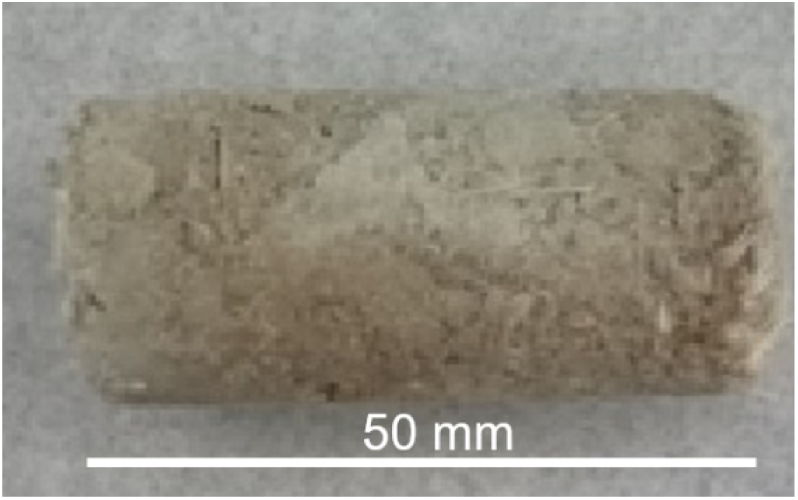
Image of bio consolidated LSS brick with *Sporosarcina pasteurii*

Compressive strength of LSS consolidated bricks was determined for five samples of each treatment. Figure 4 shows comparison of compressive strength for control samples namely, URGG00LSS, GG01LSSControl, SPSMULSS and SPGG00LSS as well as guar admixture inoculated with bacterial strain. It was found that the chemical urease treated sample gave the lowest strength of about 40 kPa. Sample containing guar gum with media and LSS without inoculation of bacterial strain (GG01LSSC) exhibited a compressive strength of 80 kPa. The samples with and without external calcium source (SPSMULSS and SPGG00LSS) showed comparable compressive strength of approximately 120 kPa. SPGG00LSS exhibiting slightly higher strength compared to SPSMULSS, where no additional calcium source was added.

**Figure 4.**
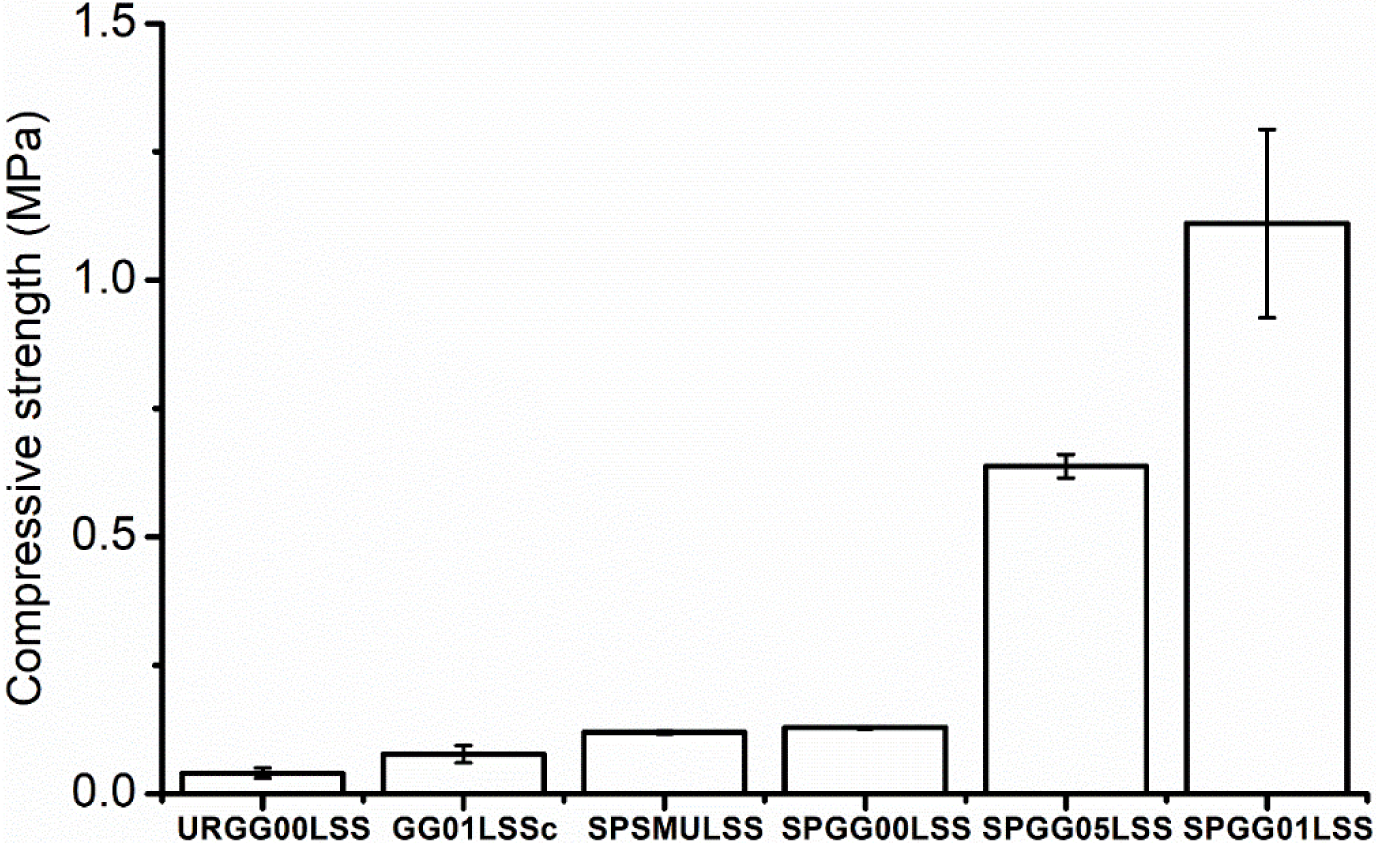
Graph showing compressive strength of bio-consolidated samples with different treatment; URGG00LSS, GG01LSSC, SPSMULSS, SPGG00LSS, SPGG05LSS & SPGG01LSS under uniaxial compression test

Compressive strength of the bio-consolidated samples with different concentration of guar exhibited seemingly brittle behavior under compression, inevitably failing via columnar fracture. In case of SP inoculated LSS soil with different percentage of guar gum admixture, the strength increased by ~400% and ~760% with 0.5% and 1% (w/w) guar gum substitution respectively. A maximum compressive strength of 1.1 MPa was observed with 1% guar gum additive. The inspiration of our work is derived from the idea of making extra-terrestrial structures, we named the bio-consolidated LSS structures as ‘space bricks’.

### B. Microstructural characterization of LSS consolidated bricks

The microstructure of LSS consolidated via bacteria induced bio-cementation was investigated by SEM and XRD techniques. The SEM image of the sample with 1% guar gum admixture (SPGG01LSS) is shown in Fig 5. It clearly depicts bio-consolidated aggregates. This aggregated dense microstructure could also be the basis for the resulting high compressive strength (cf. Fig. 4).

**Figure 5.**
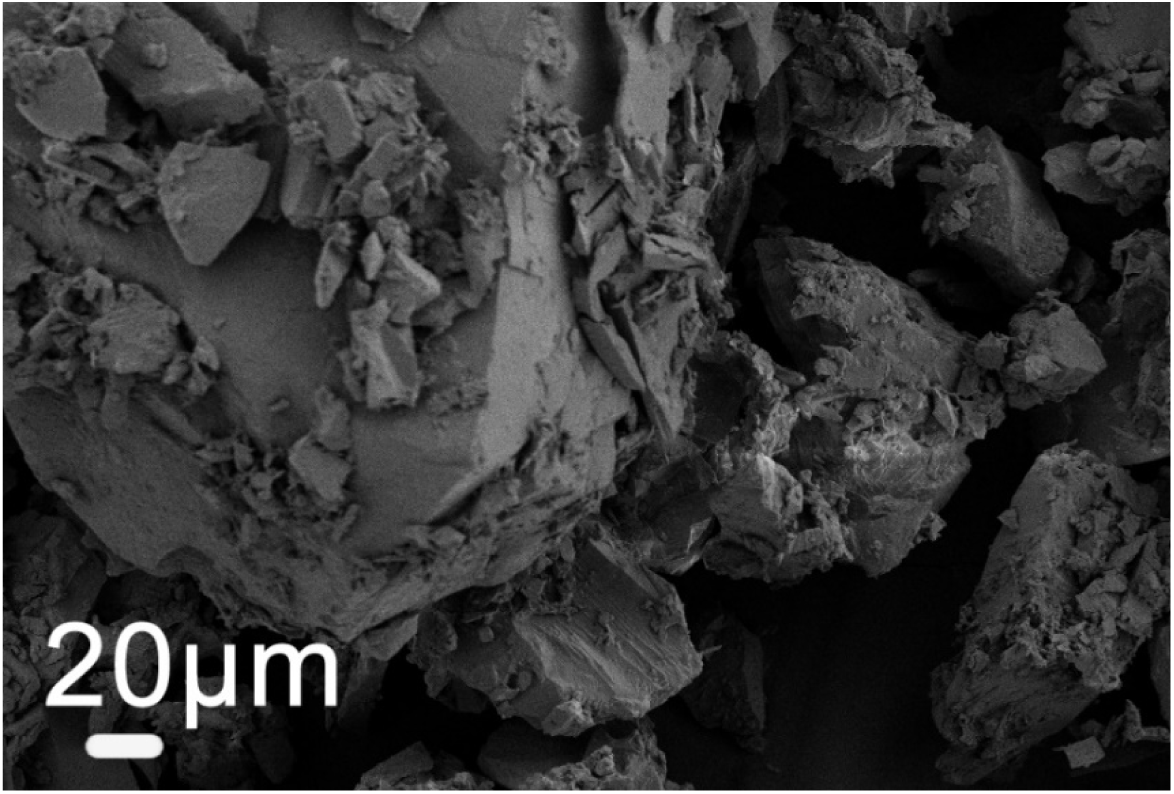
SEM images of *Sporosarcina pasteurii* mediated SPGG01LSS consolidated sample

Additional information of the structures present in the consolidated sample was obtained using XRD analysis. Figure 6 shows XRD data for both LSSC powder (control LSS sample without any treatment) and SPGG01LSS consolidated samples. Mixed phases (calcite, vaterite and aragonite) of CaCO_3_ precipitates were observed in the bio-consolidated SPGG01LSS brick sample. The peaks correspond to the calcite phase at 2θ values of 22.60°, 31.50° and matched well with ICSD File nos. 98-007-8903, 98-009-6175 while orthorhombic vaterite phase was observed in the peaks at corresponding 2θ values of 10.36°, 13.5°, 26.6°, 62.49° (ICSD File nos. 98-000-6092, 98-010-9797, 98-010-9796). Further, aragonite (ICSD File nos. – 98-011-4648, 98-011-4649) peaks was found at 2θ values of 21.85°,30.321°, 42.32°, 52.71 ° and 67.42°. Finally, Calcium carbonate peaks were also observed at 2θ values of 24.471°, 78.92° matched with ICSD File no. 98-004-0793. The other peaks besides calcium carbonate phases appear to be from the LSS matrix. On the other hand, control LSS powder sample shows Anorthosite phase along with different oxide phases as found in conventional soil samples such as silicon dioxide, aluminium oxide, calcium oxide etc. It is very important to note that the high intensity peaks correspond to the calcium carbonate phases observed in SPGG01LSS, proving that the bacteria-induced precipitation activity results in significant amounts of crystalline calcium carbonate phases. These phases could also contribute to the enhanced mechanical strength in SPGG01LSS (Fig. 4).

**Figure 6.**
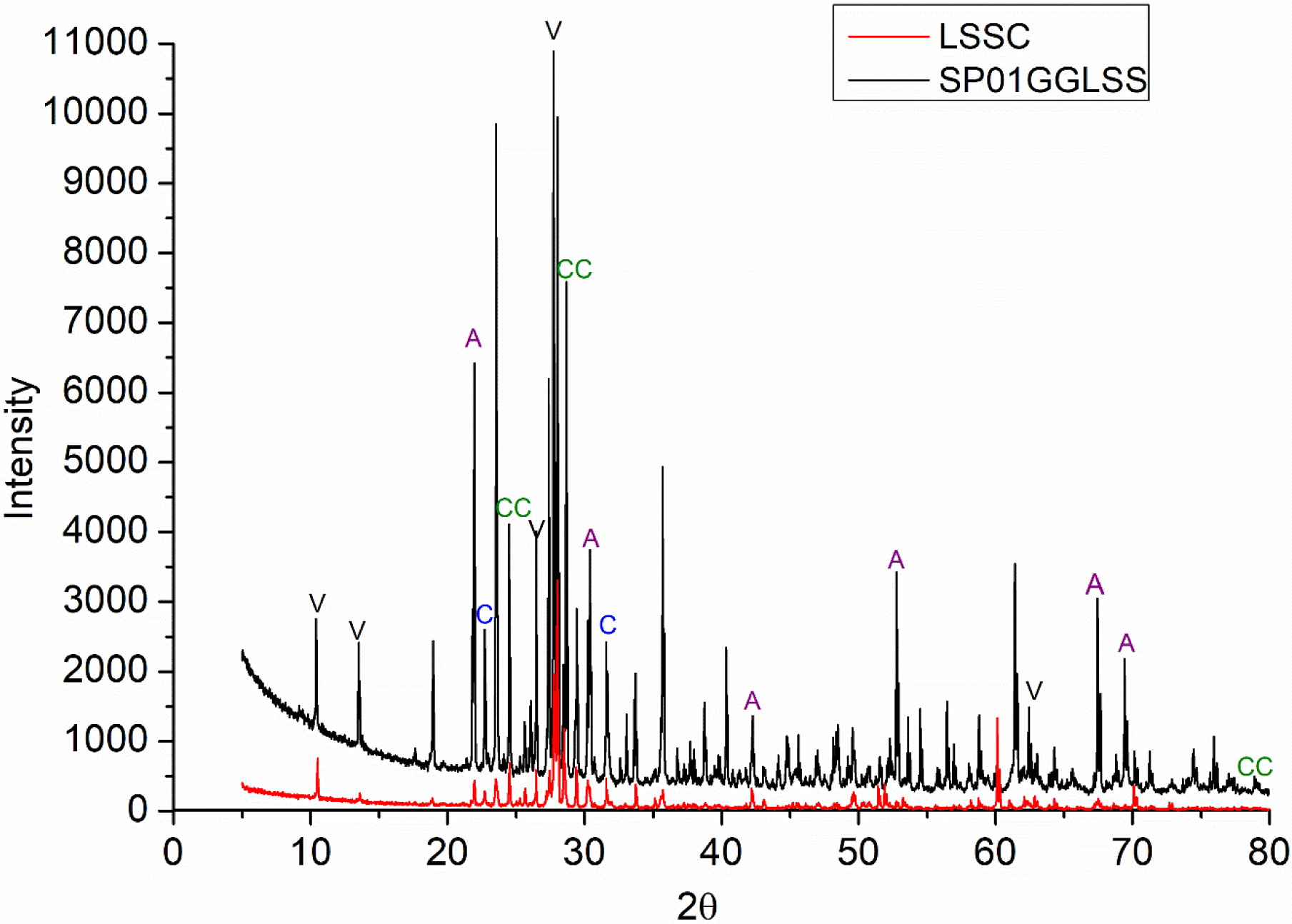
XRD spectrum of LSS consolidated bricks treated with SPGG01LSS and LSS control without any bacterial supplementation. V; vaterite, C; calcite, A; argonite, CC; calcium carbonate

### C. Experiments in Microgravity

For the bacteria-induced bio-cementation process to be useful as a potential route for in situ production of consolidated bricks with requisite strength, it is necessary to first evaluate the activity of the bacteria under space-like conditions, viz. microgravity. In order to do this, we have devised a payload design, of potential general utility in other space biology experiments using a modular approach, see the schematic Fig. 7. The design philosophy involves conceptually separating the payload into three distinct levels – the cartridge level, cassette level and the full housing. This modular approach enables changes at an individual level without affecting the other two and could potentially be used for other bio experiments where only the specific experiment (cartridge level) needs to be altered.

**Figure 7.**
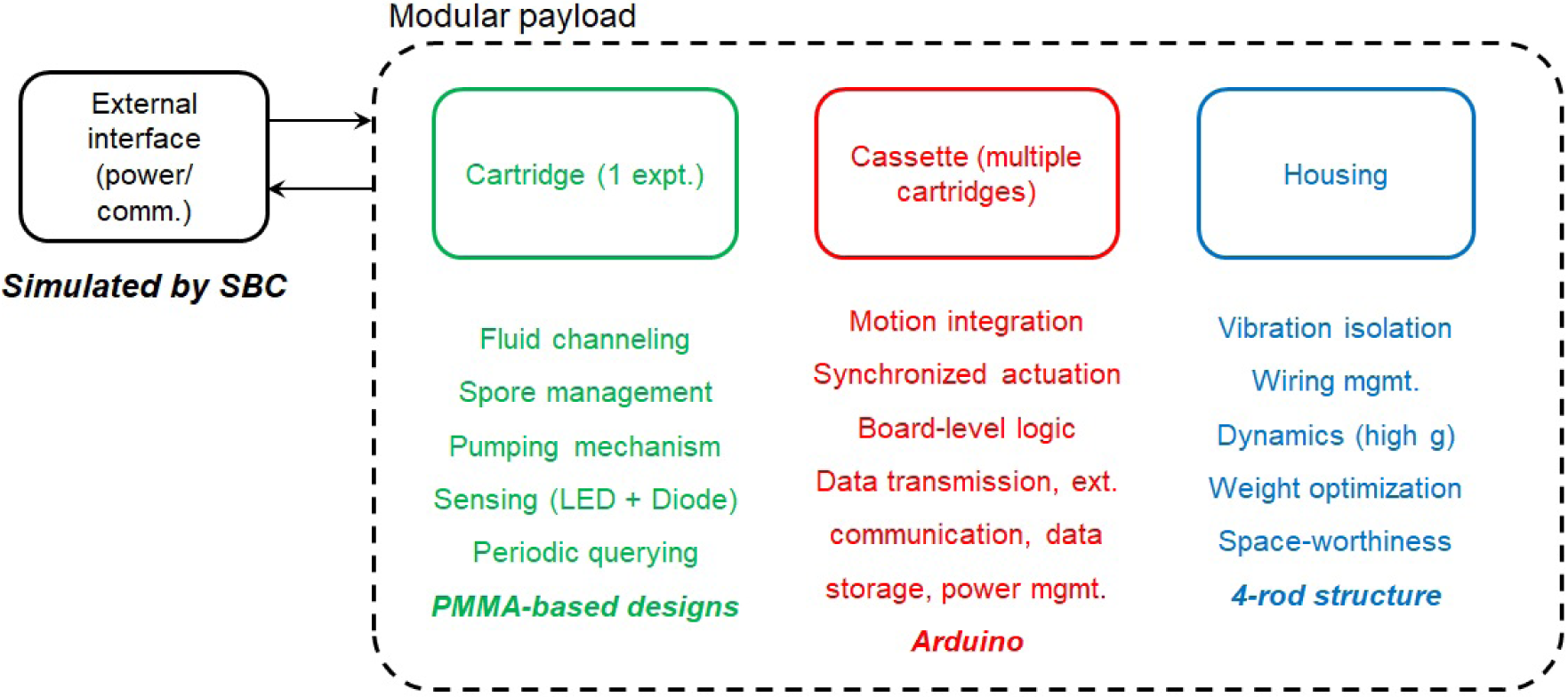
Schematic representation of modular payload design for microgravity experimentation

For the present experiment – evaluating the effect of microgravity on the activity of *Sporosarcina pasteurii* – one specific cartridge and cassette design is shown in Fig. 8. The single cartridge (Fig. 8 a) is transparent and contains a mechanism for mixing the nutrient fluid with bacteria in spore form at the start of the experiment. The mechanism is driven by a shaft that is connected to the housing (Fig. 8 b) and driven by a motor (not shown). Once bacterial growth starts, bacterial activity is measured using an LED/optical photodiode combination mounted on the periphery of the cartridge. The resulting optical density curves will be calibrated with already available ground-scale experiments. The cassette level contains 4 identical cartridges, with three independent experiments and one control (no bacteria) (Figs. 8 c and d).

**Figure 8.**
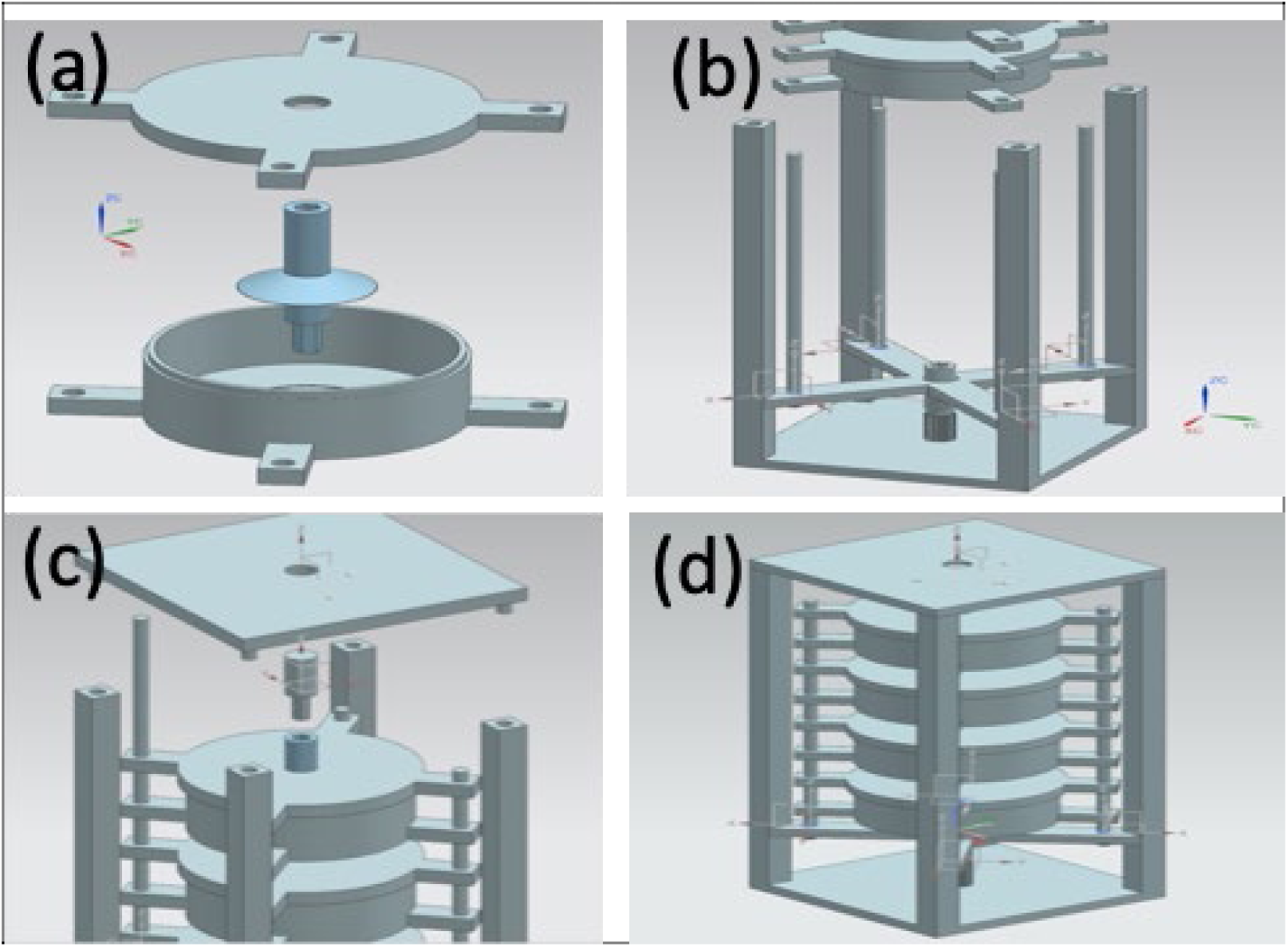
Proposed design based on the lab-on-chip’ model for microgravity experimentation

## IV Major Findings

We have shown that lunar soil simulant can be successfully bio-consolidated into a ‘space brick’ by using the MICP process. The strength of the MICP consolidated bricks was significantly increased by the use of guar gum as an additive. These results provide a strong potential for the use of MICP enabled bio-consolidation for the construction of bricks using lunar regolith for extra-terrestrial habitats. In order to further test the promise of MICP enabled bio-consolidation in extra-terrestrial conditions, it is essential to carry out tests in low-gravity conditions. Towards this end, we have proposed designs for payloads where the MICP process can be tested in microgravity conditions. Our results and proposed designs have a strong potential for utility in creating human settlements on other planets.

## V Acknowledgements

AK would like to acknowledge support from ISRO for this project.

## References

1. Taylor SL, Jakus AE, Koube KD, Ibeh AJ, Geisendorfer NR, et al. Sintering of micro-trusses created by extrusion-3D-printing of lunar regolith inks. Acta Astronaut 2018;143:1–8.

2. Wilhelm S, Curbach M. Review of possible mineral materials and production techniques for a building material on the moon. Struct Concr 2014;15:419–428.

3. F. R, J. S, H. B. Structural Design of a Lunar Habitat. J Aerosp Eng 2006;19:133–157.

4. David CW. Particle Size Distribution of Lunar Soil. J Geotech Geoenvironmental Eng 2003;129:956–959.

5. Anand M, Crawford IA, Balat-Pichelin M, Abanades S, van Westrenen W, et al. A brief review of chemical and mineralogical resources on the Moon and likely initial in situ resource utilization (ISRU) applications. Planet Space Sci 2012;74:42–48.

6. Vamsi KB. First demonstration on direct laser fabrication of lunar regolith parts. Rapid Prototyp J 2012;18:451–457.

7. Crockett RS, Fabes BD, Nakamura T, Senior CL. Construction of large lunar structures by fusion welding of sintered regolith. In: Engineering, construction, and operations in space IV. ASCE; 1994. pp. 1116–1127.

8. A. TL, T. MT. Microwave Sintering of Lunar Soil: Properties, Theory, and Practice. J Aerosp Eng 2005;18:188–196.

9. Anbu P, Kang C-H, Shin Y-J, So J-S. Formations of calcium carbonate minerals by bacteria and its multiple applications. Springerplus 2016;5:250.

10. Hammes F, Verstraete* W. Key roles of pH and calcium metabolism in microbial carbonate precipitation. Rev Environ Sci Biotechnol 2002;1:3–7.

11. Zhu T, Dittrich M. Carbonate Precipitation through Microbial Activities in Natural Environment, and Their Potential in Biotechnology: A Review. Frontiers in Bioengineering and Biotechnology 2016;4:4.

12. Ghosh T, Bhaduri S, Montemagno C, Kumar A. *Sporosarcina pasteurii* can form nanoscale calcium carbonate crystals on cell surface. PLoS One 2019;14:e0210339.

13. Achal V, Pan X. Characterization of Urease and Carbonic Anhydrase Producing Bacteria and Their Role in Calcite Precipitation. Curr Microbiol 2011;62:894–902.

14. Vahabi A, Ramezanianpour AA, Sharafi H, Zahiri HS, Vali H, et al. Calcium carbonate precipitation by strain *Bacillus licheniformis* AK01, newly isolated from loamy soil: a promising alternative for sealing cement-based materials. J Basic Microbiol 2015;55:105–111.

15. Bhaduri S, Debnath N, Mitra S, Liu Y, Kumar A. Microbiologically induced calcite precipitation mediated by *Sporosarcina pasteurii*. JoVE (Journal Vis Exp 2016;e53253.

16. Lambert SE, Randall DG. Manufacturing bio-bricks using microbial induced calcium carbonate precipitation and human urine. Water Res 2019;160:158–166.

17. Randall DG, Naidoo V. Urine: The liquid gold of wastewater. J Environ Chem Eng 2018;6:2627–2635.

18. Muguda S, Booth SJ, Hughes PN, Augarde CE, Perlot C, et al. Mechanical properties of biopolymer-stabilised soil-based construction materials. Géotechnique Lett 2017;7:309–314.

19. Aminpour M, O’Kelly BC. Applications of biopolymers in dam construction and operation activities. In: Proceedings of the 2nd International Dam World Conference, Lisbon, Portugal. Laboratório Nacional de Engenharia Civil, Lisbon, Portugal; 2015. pp. 937–946.

20. Chang I, Im J, Prasidhi AK, Cho G-C. Effects of Xanthan gum biopolymer on soil strengthening. Constr Build Mater 2015;74:65–72.

21. M. Annadurai, I. Venugopal and K. Muthukkumaran. Road map to build civil engineering structures on the moon-an overview10th IAA symposium on the future of space exploration: towards the moon village and beyond. In international academy of astronautics (IAA), Torino, Italy, June 27-29, 2017.

